# Untargeted metabolomics reveals PTI-associated metabolites in tomato

**DOI:** 10.1101/2023.06.15.544816

**Authors:** Lina Muñoz Hoyos, Petra Anisha Wan, Chen Meng, Karin Kleigrewe, Corinna Dawid, Ralph Hückelhoven, Remco Stam

## Abstract

Plants employ a multi-layered innate immune system to detect and fend off invading fungal pathogens. In one such layer, recognition of Pathogen- or Microbe-Associated Molecular Patterns or elicitors, triggers a signaling cascade that leads to defence against the pathogen and ultimately Pattern-Triggered Immunity (PTI). Secondary Metabolites (SMs) are expected to play an important role in this kind of resistance, because they are potentially mycotoxic compounds. Tomato plants inoculated with *Alternaria solani* show clear symptoms of infection 5 days after inoculation. Whereas plants inoculated with *Alternaria alternata* remain symptomless. We hypothesized that pattern-triggered induction of resistance-related metabolites in *Solanum lycopersicum* contribute to the resistance against *A. alternata*, yet such SMs are suppressed in a compatible interaction. We compared the metabolomic profile (metabolome) of *S. lycopersicum* at two time points (3 and 24 hours) after treatments with *A. alternata, A. solani* and the fungal elicitor chitin and identified SMs that are involved in the early defence response of tomato plants. Our study revealed differential metabolome fingerprints and shows that the molecular composition of *A. alternata* and chitin-induced indeed show larger overlap with each other than with the *A. solani-*induced metabolome. We identify 65 candidate metabolites possibly associated with pattern-triggered resistance in tomato plants, including the alkaloid, trigonelline, for which we can confirm that it inhibits fungal growth *in vitro* when supplied at physiological concentrations. Our findings show that a true, pattern-triggered, chemical defence is mounted against *A. alternata* and that it contains mycotoxin compounds previously unidentified in tomato, that could be interesting for future crop protection strategies.

## Introduction

Tomato is one of the most important vegetable crops worldwide, with a total fresh production that exceeded 187 million tons in 2020 (FAOSTAT, 2020). It can be grown in a wide range of climates from tropical to temperate and it can also be cultivated under cover conditions when outdoor temperatures are not favorable. Because of its wide use and nutritional values, there is a high demand for both fresh market and processed tomato varieties. However, a diversity of bacterial, viral, nematode, and fungal diseases, have made it difficult to grow tomatoes for commercial purposes (Bozbuga et al., 2022).

Due to intensive selection and inbreeding through domestication, the cultivated tomato has a limited genetic variety; as a result, they are more vulnerable to disease epidemics (Schauer et al., 2008). Early blight (EB) is one of the most common diseases of tomato caused by several species of *Alternaria* fungi, predominantly *A. solani* (Adhikari et al., 2017). The typical disease symptoms are dark brown to black lesions with concentric rings on the leaves, which result in leaf browning and leaf drop in severe cases, these lesions can occur over a wide range of environmental conditions causing severe reductions (35–78%) in yield (Özer Çalış, 2011). During the infection process, *A. solani* produces germ tubes to penetrate host tissues or directly invades from stomata or wounds adjacent to epidermal cells. It secretes metabolites to break down host cell inclusions, thereby causing infection (Adhikari et al., 2017). Complete resistance of tomato against *A. solani* has not been observed (Chaerani et al., 2007; Ray et al., 2015). Notably, some tomato genotypes are resistant to *A. alternata*, another causal agent of EB diseases (Moghaddam et al., 2022). However, whereas weak correlations can be found between several morphological properties and *A. alternata* resistance, it remains unclear which specific biochemical properties are responsible for the resistance of the genotypes. Another study evaluating tomato genotypes for early blight disease caused by different *Alternaria* species, did not find any genotype that were resistant to *A. alternata* isolates tested (Akhtar et al., 2019). Overall, while some tomato genotypes may show resistance to some *Alternaria* species or isolates, the mechanism underlying this resistance is not well understood yet.

Pathogens use different strategies to inhibit constitutive and induced plant defences, including degradation of preformed antimicrobial compounds and the production of molecules that suppress induced plant defences. Necrotrophic pathogens like *A. solani* colonize the cell through production of cell wall-degrading enzymes and phytotoxins killing them and acquire nutrients from the dead plant cells (Chaerani & Voorrips, 2006; Sadeghi et al., 2022). Plants are able to perceive these pathogens or their Pathogen or Microbe Associated Molecular Patterns (PAMPs or MAMPs, or patterns in short), also referred to as elicitors, such as the cell-wall component chitin, with specific receptors that trigger the mitogen-activated protein kinase (MAPK) cascades and activating hormone (jasmonates and ethylene)-dependent and hormone-independent signaling, which facilitates the mounting of a defence response against these necrotrophs and results in Pattern Triggered Immunity (PTI). This response involves the activation of specific transcription factors that result in the production of antifungal proteins or accumulation of defensive secondary metabolites (phytoalexins) (Muñoz-Hoyos & Stam, 2023; Pandey et al., 2016). The three main groups of secondary metabolite products in plants are terpenes, phenolics, and nitrogen-containing compounds (Taiz & Zeiger, 1991). Cultivated tomato, *S. lycopersicum* and its wild relative in this genus contain multiple representatives of these three major groups (Duffey & Stout, 1996). In addition, tomatoes and wild tomatoes produce methyl ketones and acylsugars, which are present in trichomes and play an important role in resistance to some microorganisms (Glas et al., 2012) and the presence of flavonoids, alkaloids, saponins, tannins, terpenoids, glycosides and steroids in plant extracts reduced the growth of *A. solani in vitro* (Ahmad et al., 2017). Like most of the members of the family *Solanaceae*, tomatoes also contain alkaloids. A well-known steroidal glycoalkaloid and saponin constitutively produced in tomato plants is *α*-tomatine. In healthy tomato leaf tissue, *α*-tomatine exists at sufficient concentrations to inhibit the growth of many fungi *in vitro* (Roddick, 1974) and it is therefore considered the major preformed compound that protects plant against attack by wide range of potential fungal pathogens (Osbourn, 1996a). Fungal tomato pathogens are resistant to *α*-tomatine *in vitro* (Sandrock & VanEtten, 1998) because of their capability to detoxify this compound (Oka et al., 2006). Pathogens, such as *A. solani, Botrytis cinerea* and *Fusarium oxysporum* s. sp. *Lycopersici* are known to produce extracellular enzymes that hydrolyze sugars from α-tomatine in various ways to be able to infect the plant (Roldán-Arjona et al., 1999). Seeing the important roles that secondary metabolites play in defence against different pathogens, we hypothesize that in addition to *α*-tomatine, induced compounds must be synthesized by tomato plants to confer resistance to *Alternaria* species, and that elicitors such as chitin can trigger the production of these defence compounds.

Omics-based tools, such as untargeted metabolomics are potent techniques to investigate molecular changes operating during plant-pathogen interaction. Untargeted metabolomics involves global metabolic profiling with high throughput technology such as liquid chromatography coupled to tandem mass spectrometry (LC-MS/MS), followed by identification, quantification and visualization of generated metabolites (Allwood et al., 2021; Muñoz-Hoyos & Stam, 2023). By analyzing the metabolites present in plants that are resistant or susceptible to a particular stressor, biomarkers or patterns can be identified to develop strategies for improving plant resistance to different pathogens. In this study, we aim to identify pattern-triggered bio-active compounds that play a role in the resistance of cultivated tomato plants to *A. alternata* and validate their biological effects on the pathogen *in vitro*. For this, we perform untargeted metabolomics analysis with tomato leaves at three hours and 24 hours after infection with *A. alternata*, *A. solani* and elicitation with chitin, representing the times in which the fungal spores germinate and try to penetrate the plant to establish disease. The identification of compounds as biochemical markers may be useful as screening tools for plant material in breeding programs.

## Materials and methods

### Plant material

Seeds of *S. lycopersicum* (HEINZ 1706) were obtained from the Centre of Genetic Resources, the Netherlands (CGN). To assure homogenic material, cuttings were made of a single mother plant after 6 weeks of sowing. Three week old cuttings were used for the metabolomics comparison. Plants were grown in controlled chamber conditions (12 h light, 22-24 °C).

### Fungal isolates

*A. alternata* was isolated from the leaves of the wild tomato plant (*Solanum chilense*) showing signs of leaf spot (Schmey et al., 2022). *A. solani* (1117-1) was isolated from tomato in Freising (Germany). The identity of the fungal isolates was confirmed based on morphological (Simmons, 2007) and molecular features (Schmey et al., 2022).

### Pathogen inoculation, plant elicitation and plant infection assays

*A. alternata* and *A. solani* strains were cultured on synthetic, nutrient-poor agar (SNA) plates at 25°C, 12UV-A light, 12 h darkness and 85% humidity for ten days. The spores were harvested by scraping them with an inoculation loop from the plates and placing in water. The spore concentration was determined under the microscope with a Thoma counting chamber, and the suspension was diluted to a concentration of 3×10^4^ spores per mL. Chitin derived from shrimp shell (C9752; Sigma-Aldrich) was ground and re-suspended in water, a final concentration of 50 µg/mL was used for experiments. Spray inoculation was performed for elicitation and infection of the plants (5 mL per plant). 24 plants of *S. lycopersicum* were sprayed with *A. alternata*, *A. solani*, chitin and water and the leaves were collected after 3 and 24 hours post inoculation [hpi], (3 plants per treatment and time point). Similar samples were used for both chlorophyll measurements and for untargeted MS. The leaves from every plant (3 replicates) were collected and immediately frozen in liquid nitrogen. Samples were homogenized using liquid nitrogen and porcelain mortars and pestles. Grounded samples were freeze-dried and 100 mg of grounded sample were used for extraction. Each of the samples was extracted using 1,000 µL of extraction solution (90% methanol, 10% water) shaking in a vortex for 30 min. Samples were then centrifuged for 15 min at 12,000 rpm and 200 µL of the supernatant was collected into glass tubes and stored at -20°C.

### Red light chlorophyll fluorescence measurement for cell death quantification

Red light emission of chlorophyll fluorescence can be measured for quantification of cell death (Landeo Villanueva et al., 2021). Leaf discs (Ø 4mm) of *S. lycopersicum* treated with *A. solani*, *A. alternata* or water were floated on water in black 96-well plates and chlorophyll fluorescence was measured with a plate reader (Tecan Infinite F200 PRO, excitation 535 nm, emission 590nm, 25 flashes, integration time 20µs, 4×4 reads per well, gain set to 80) as relative fluorescence units (RFU). Values of all reads per well were summed up.

### Cultivation of fungal isolates for MS of growth media

The *Alternaria* isolates were cultivated in *S. lycopersicum* host broth medium to obtain the exometabolome (pool of exogenous metabolites). 20 g of *S. lycopersicum* leaves were boiled with 500 mL of distilled water for 15 minutes, the leaves were then filtered using a stainless-steel mesh strainer. 150 mL of the liquid medium was transferred into 3 different polycarbonate Erlenmeyer flasks, the pH was adjusted to 7 using formic acid and was later autoclaved at 121 °C for 20 min. The sterile liquid media was inoculated with 25 µL of the spores suspension of *A. alternata* (8.75 x 10^5^ spores per milliliter) and 100 µL of the spores suspension of *A. solani* (2×10^5^ spores per milliliter) to receive equal amount of total spores. The fungi were cultivated in the dark (25 °C, 100 rpm) and the isolates were exposed to artificial light for half an hour a day. After 11 days of cultivation the liquid medium was filtered with filter paper Whatman No.1 to separate the mycelium. 5 mL of the filtrate were pipetted and transferred to 15 mL Falcon tubes and stored at -20°C. 3 different types of samples were measured: “*A. alternata* exo” (liquid broth media from *S. lycopersicum* after filtering *A. alternata* mycelium), “*A. solani* exo” (liquid broth media from *S. lyopersicum* after filtering *A. solani* mycelium) and “control” (liquid broth media from *S. lycopersicum* without fungus inoculum). 200 µL of the liquid media were used for extraction. Each of the samples was extracted using 1,000 µL of extraction solution (90% methanol, 10% water) shaking in a vortex mixer for 30 min. Samples were then centrifuged for 15 min at 12,000 rpm and 200 µL of the supernatant was collected into glass tubes and stored at -20°C.

### Metabolomics analysis

The untargeted metabolite analysis was performed using a Nexera UHPLC system (Shimadzu, Duisburg, Germany) coupled to a Q-TOF mass spectrometer (TripleTOF 6600, AB Sciex, Darmstadt, Germany). Separation of the samples was performed using a UPLC BEH Amide 2.1 × 100 mm, 1.7 µm analytic column (Waters, Eschborn, Germany) with a 400 µL/min flow rate. The mobile phase was 5 mM ammonium acetate in water (eluent A) and 5 mM ammonium acetate in acetonitrile/water (95/5, v/v) (eluent B). The gradient profile was 100% B from 0 to 1.5 min, 60% B at 8 min and 20% B at 10 min to 11.5 min and 100% B at 12 to 15 min. A volume of 5 µL per sample was injected. The autosampler was cooled to 10 °C and the column oven heated to 40 °C. Every tenth run a quality control (QC) sample, which was pooled from all samples, was injected. The samples were measured in a randomized order and in the Information Dependent Acquisition (IDA) mode. MS settings in the positive mode were as follows: Gas 1 55 psi, Gas 2 65 psi, Curtain gas 35 psi, Temperature 500 °C, Ion Spray Voltage 5500 V, declustering potential 80 V. The mass range of the TOF MS and MS/MS scans were 50–2000 *m/z* and the collision energy was ramped from 15–55 V. MS settings in the negative mode were as follows: Gas 1 55 psi, Gas 2 65 psi, Cur 35 psi, Temperature 500 °C, Ion Spray Voltage –4500 V, declustering potential –80. The mass range of the TOF MS and MS/MS scans were 50–2000 *m/z* and the collision energy was ramped from -15 – -55 V. The “msconvert” from ProteoWizard was used to convert raw files to mzXML (de-noised by centroid peaks). The bioconductor/R package xcms was used for data processing and feature identification (Smith et al., 2006). More specifically, the matched filter algorithm was used to identify peaks (full width at half maximum set to 7.5 s). Then the peaks were grouped into features using the “peak density” method. The area under the peaks was integrated to represent the abundance of features. The retention time was adjusted based on the peak groups presented in most of the samples (Benton et al., 2010; Tautenhahn et al., 2008). To annotate possible metabolites to identified features, the exact mass and MS/MS fragmentation pattern of the measured features were compared to the records in HMBD (Wishart et al., 2007) and the public MS/MS database in MSDIAL, referred to as MS1 and MS2 annotation, respectively. The resulting peak list was uploaded into MetaboAnalyst 5.0 (Xia et al., 2009), a web-based tool for metabolomics data processing, statistical analysis, and functional interpretation where statistical analysis and modeling were performed. Missing values were replaced using a (K-nearest neighbor) KNN missing value estimation. Data filtering was implemented by detecting and removing non-informative variables that are characterized by near-constant values throughout the experimental conditions by comparing their robust estimate interquartile ranges (IQRs). Data were autoscaled. Out of 4496 mass features originally detected, 2499 were used for the principal least square discriminant analysis (PLS-DA) (Lê Cao et al., 2011). For the identification of candidate metabolites, the individual mass features that contributed to the separation between the different treatments were further characterized by applying a range of univariate and multivariate statistical tests to determine the importance including PLS-DA importance variables, t-test, and random forest. For identity identification of the candidate compounds SIRIUS 4 was used (Dührkop et al., 2019). The data is available through MassIVE MSV000092161.

### Bioassay on the antifungal effect of trigonelline on the growth of Alternaria

Physiological concentrations of trigonelline in tomato leaves were estimated based on literature (328 ppm) (Tyihák et al., 1988). *Alternaria* isolates were grown on SNA plates supplemented with different concentrations of trigonelline or nicotinic acid spanning both sides of the previously mentioned value (0.0328 mg/mL, 0.0656 mg/mL, 0.328 mg/mL, 1.64 mg/mL, 3.28 mg/mL). The fungi were grown at 25 °C, 12UV-A light, 12 h darkness and 85% humidity. For growth rate assays, radial growth was measured at regular intervals (1, 2, 3, 4, 5, 7, 9 and 14 days post inoculation) after placing a 4 mm agar plug in the center of the petri dishes. The spore germination was assessed microscopically. For each treatment and time point 50 spores were observed.

## Results

### S. lycopersicum (Heinz 1706) is resistant to A. alternata (CS046)

Prior investigation into whether the secondary metabolites differ between our treatments, e.g. elicitors, and compatible or incompatible pathogens, we performed infection assays on *S. lycopersicum* plants inoculated with *A. solani* and *A. alternata*. To confirm the infection phenotype, we studied a subset of leaves for up to 4 dpi. We visualized the necrotic lesions by measuring chlorophyll fluorescence (Fig. 1A), which is associated with cell death (Landeo Villanueva et al., 2021). We observed necrotic lesions in plants treated with *A. solani*, whereas the plants treated with *A. alternata* did not show any symptoms (Fig. 1B). In the same manner, leaves treated with *A. solani* showed increased chlorophyll fluorescence compared with *A. alternata* and mock treatments, validating the ability of our *A. solani* isolate to infect *S. lycopersicum* plants in contrast to our *A. alternata* isolate.

**Figure 1.**
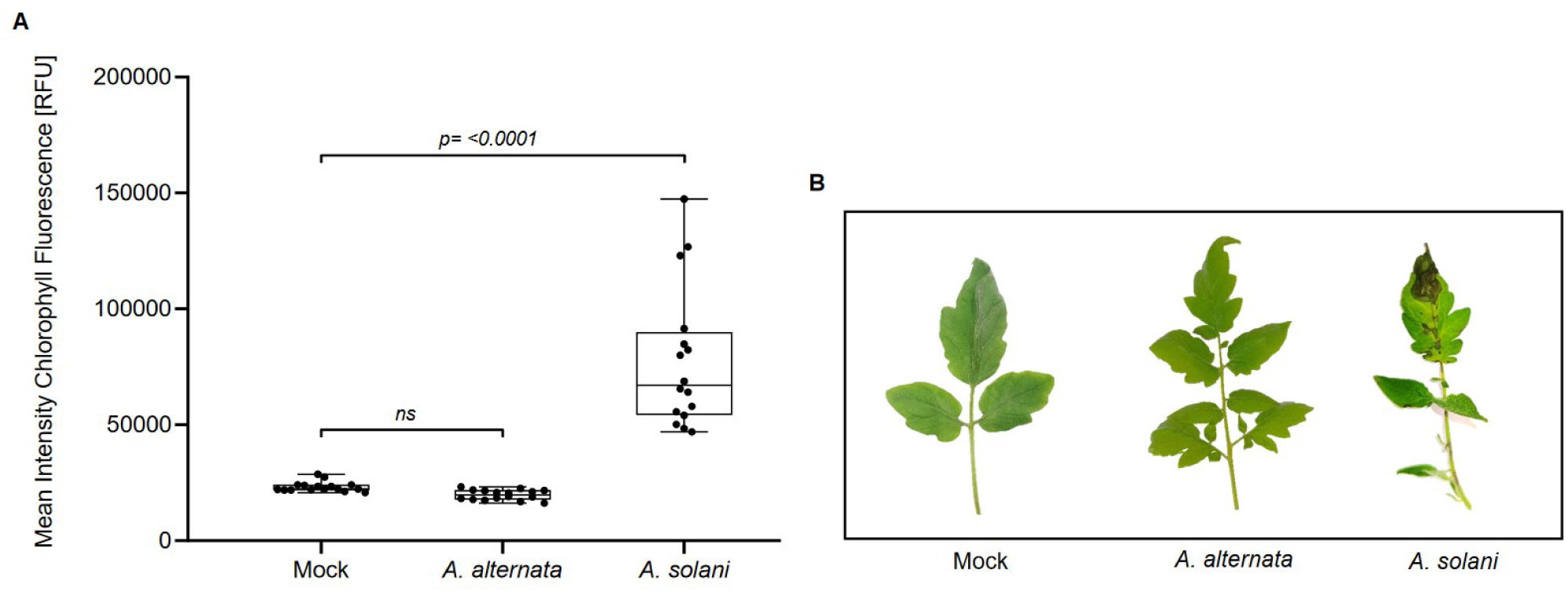
**(A)** Chlorophyll fluorescence measurement serves as proxy for cell death upon infection of tomato plants inoculated with *A. alternata* or *A. solani*. n= 16 leaf discs. Y-axis show fluorescence in relative fluoresence units (RFU). **(B)** Tomato leaves after 4 days of spray inoculation with *A. alternata*, *A. solani* and water (Mock).

#### Both fungi can break down α-tomatine

It is known that *α*-tomatine is an important phytoanticipin against fungi that needs to be detoxified prior to successful infection (Oka et al., 2006; Ökmen et al., 2013). To assess the presence of phytoanticipins that could inhibit the growth of *A. alternata* on tomato, we performed inoculation in tomato leaf broth medium, hypothesizing that this medium contains all phytoanticipins, but that it cannot show induced defence responses. Both fungi are able to grow in this medium, confirming that phytoanticipins do not play a major role in the resistance against *A. alternata* (Fig 2A). To corroborate these results and to show that both fungi are able to detoxify *α*-tomatine, we performed untargeted metabolomics on the Heinz exometabolome (exo) (*S. lycopersium* broth liquid medium after infection with the fungi and the mycelium being filtered out) and Heinz control medium (*S. lycopersicum* broth liquid medium without fungal inoculum). We then looked specifically for features that can be associated with typical known phytoanticipins, such as *α*-tomatine. We found that *A. solani* exo and *A. alternata* exo showed a significant decrease in the amount of *α*-tomatine compared with the control (Fig 2B) confirming a possible degradation or detoxification of this compound into other molecules. To confirm degradation or detoxification of *α*-tomatine, we looked at presence of known related compounds. Tomatidine, a glycoalkaloid and precursor and possible degradation product of *α*-tomatine (Oka et al., 2006), significantly accumulated in *A. alternata* exo compared with the control, but not in *A. solani exo* samples. We found a similar pattern for *β*1-tomatine, a compound known to be the result of *α*-tomatine detoxification by fungi such as *B. cinerea* (Quidde et al., 1998), where an increase in accumulation levels was observed in *A. alternata* exo compared with *A. solani* exo samples. On the other hand, (23*R*)-23-acetoxytomatidine, a steroidal alkaloid, strongly accumulated in *A. solani exo* compared with *A. alternata exo* and the control. This suggests that both fungi are able to degrade (different) phytoanticipins, but use different strategies for its degradation.

**Figure 2.**
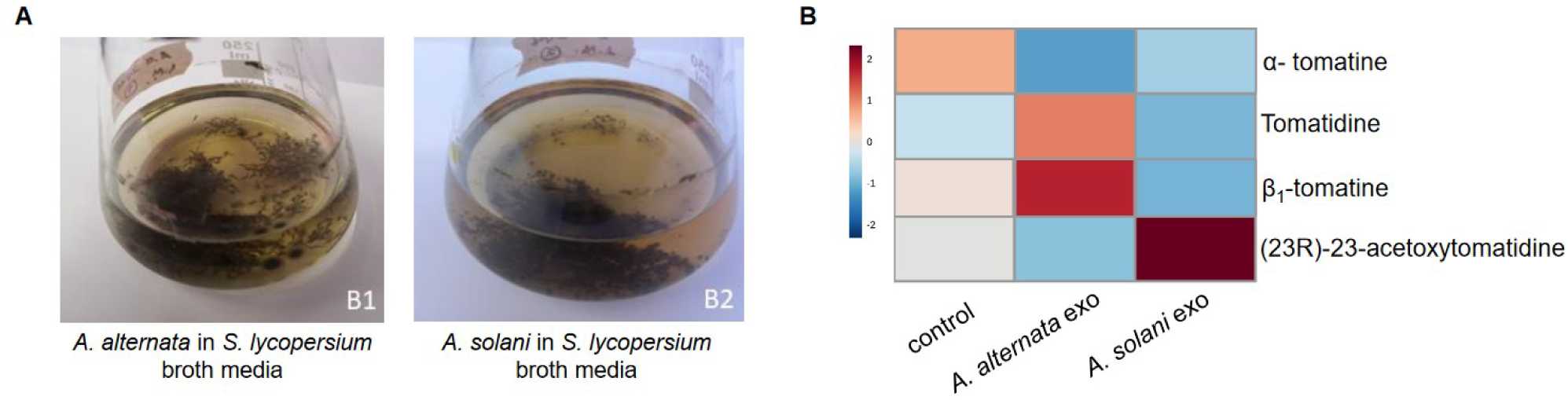
**(A)** *S. lycopersicum* broth liquid media after 11 days of inoculation with *A. alternata* and *A. solani* **(B)** Heatmap representing the relative levels of *α*-tomatine and derivatives in the control, *A. alternata* exo and *A. solani* exo samples. Values range from -2 (dark blue) to 2 (dark red).

#### Metabolic profiles differ after treatment with *A. alternata*, *A. solani* and chitin

To identify PAMP and pathogen-associated metabolome changes in tomato plants, we performed untargeted metabolomics analysis in tomato leaves treated with *A. alternata*, *A. solani* and chitin at two different times (3 and 24 hpi). Subsequently, we conducted a multivariate analysis to compare the changes in the metabolite profiles for the different treatments. The principal component analysis (PCA) score plot for all the treatments revealed two separated groups corresponding to samples treated with *A. alternata*, *A. solani,* chitin, and samples treated with water (mock), suggesting a possible large generic “stress” response during treatment (Fig S1). To visualize the smaller differences, we created PLS-DA score plots for the plants treated with *A. alternata*, chitin and *A. solani* compared with mock, at two different time points (3 and 24 hpi). These show that component 1 accounts for 20.5%, 22.7%, 17.6% and component 2 accounts for 6.4%, 3.5%, 9.2% of the variance, respectively (Fig 3 A, B, C). The samples that overlapped in the PCA score plot (*A. alternata*, chitin and *A. solani* treatments) are clearly differentiated in the PLS-DA score. The differentiation of 3 and 24 hpi samples is also apparent. We conclude that there is a significant reprogramming of the metabolome in the leaves following the different treatments.

**Figure 3.**
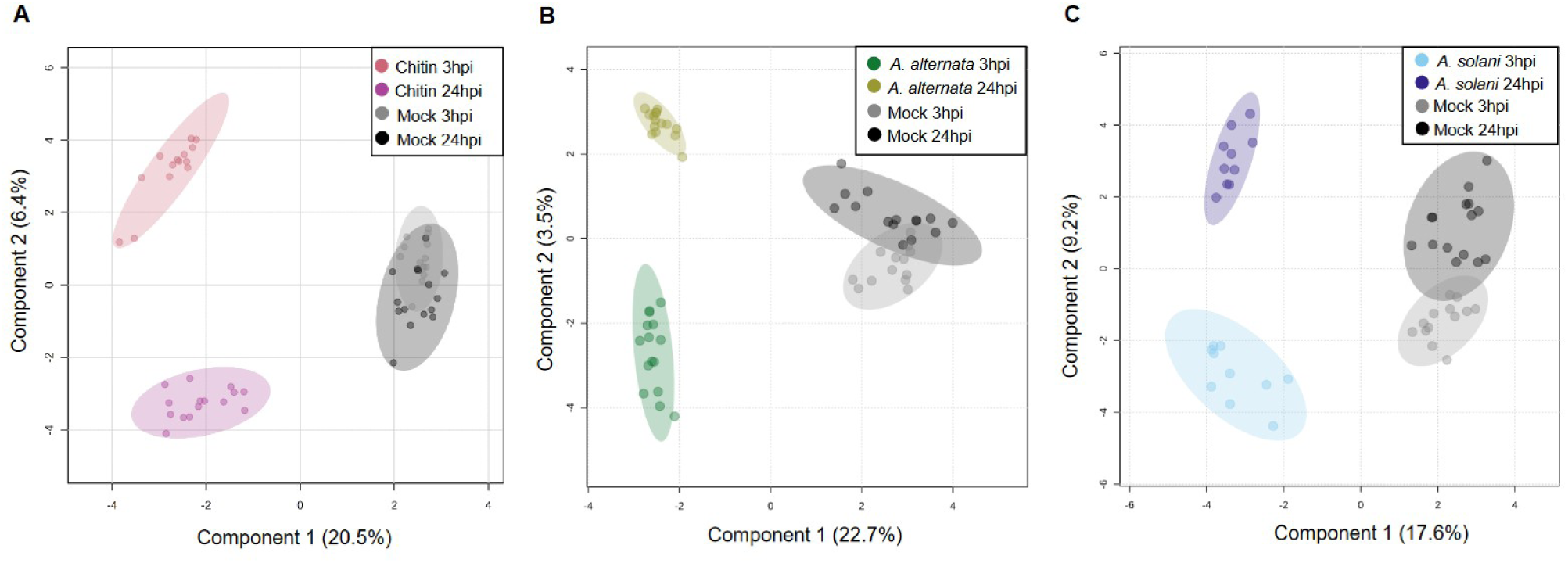
**(A)** PLS representation of statistical analysis for leaves treated with chitin, samples marked in pink and dark magenta **(B)** PLS representation of the statistical analysis for leaves treated with *A. alternata,* samples marked in dark yellow and green **(C)** PLS representation for *A. solani*, samples marked in soft blue and dark blue. All the different treatments were compared with mock, samples marked in grey and black. Sample size for every treatment (*n=15*). Confidence regions of 95% marked in circles for each group. Axes show the first (x axis) and second (y axis) components. The percentage variance explained is indicated on the axes.

### A. alternata, A. solani and chitin treatments induced a SM response

To look for the number of features that significantly accumulated or were reduced in each treatment 3 and 24 hpi, we performed volcano diagrams analysis (FDR adjusted *P*-value < 0.05 and FC > 1). *A. alternata*, *A. solani* and chitin treated samples were compared with mock samples independently to determine the differentially abundant features between all the treatments. This yields a total of 622 and 748, 606 and 618, 413 and 502 mass spectrometric features that were significantly accumulated for *A. alternata*, chitin and *A. solani* at 3 and 24 hpi respectively (Supplementary Figure 2A, B). In most cases, the number of features that accumulated to significantly higher levels were higher than those that were reduced in abundance. For all the different treatments, a greater number of features changed in relative abundance at 24 hpi compared to 3 hpi. This indicates a clear initial response and a change of the metabolome after all the treatments. We observed the highest number of differential features in *A. alternata* treated plants, implying that a successful defence leads to a stronger reprogramming towards accumulation of SMs than chitin treatment or successful infection.

Our hypothesis is that resistance of tomato to *A. alternata* may be pattern triggered immunity and that this can be seen in the accumulation of secondary metabolites that are active against the pathogen. If accurate, *A. alternata*-triggered secondary metabolites alterations in defence should overlap with those caused by the chitin treatment. We looked for the significantly accumulated features that both treatments shared. The number of unique differentially abundant mass spectrometric features observed for each treatment after 3 and 24 hpi was higher in plants treated with chitin compared to plants treated with *A. alternata* and *A. solani* suggesting that chitin triggers a more generic response (Fig 4A, B). The amount of features that overlaps between chitin and *A. alternata* treatments is higher compared to that one shared between chitin and *A. solani* treatments at 3 hpi and 24 hpi for positive and negative ionization. Finally, despite successful infection, the samples treated with *A. solani* exhibited the lowest amount of features compared to the other treatments. These results suggest that, indeed, resistance to *A. alternata* is largely pattern-triggered and furthermore that the successful infection of *A. solani* in *S. lycopersicum* limits differential accumulation of metabolites.

**Figure 4.**
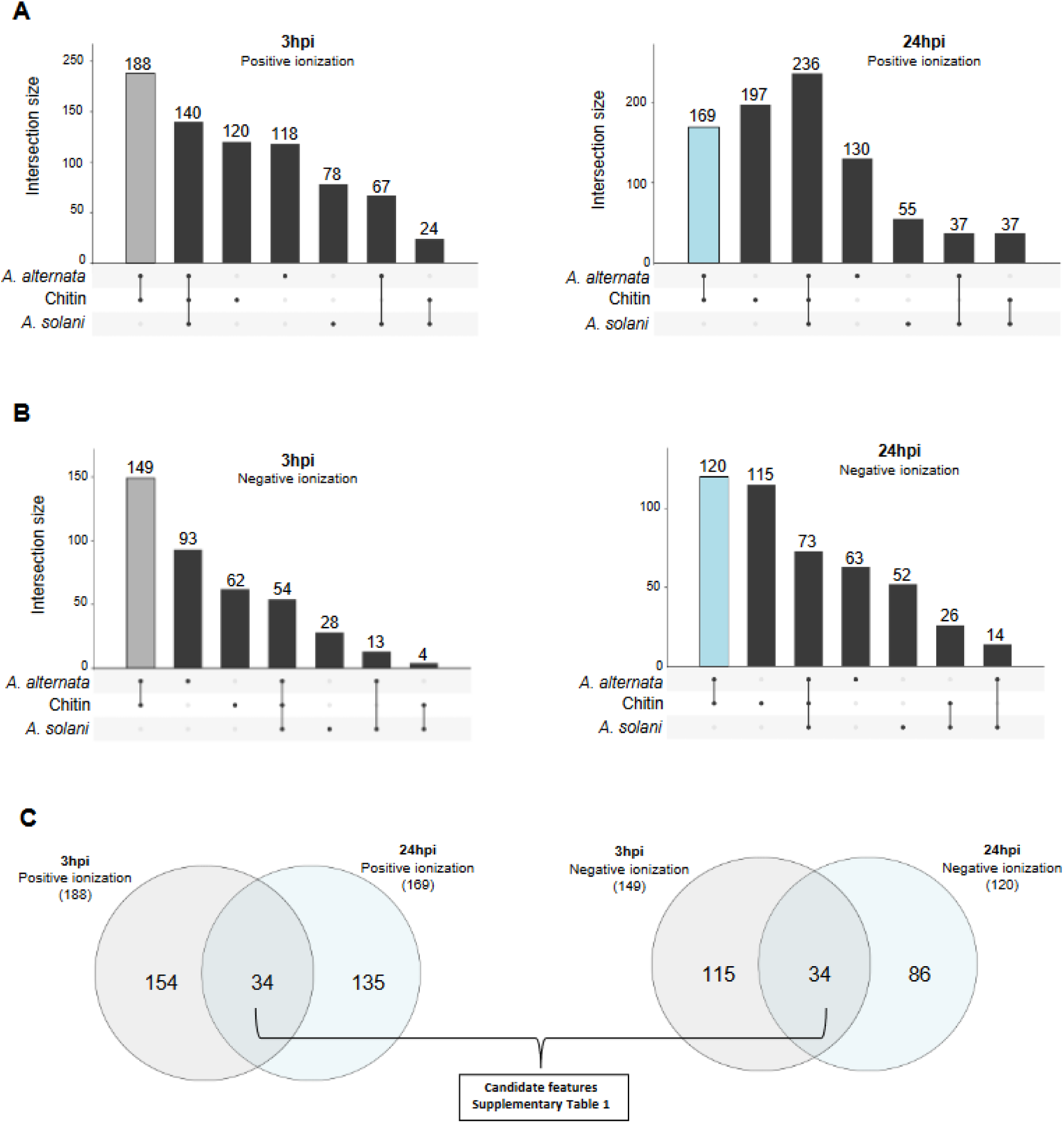
**(A)** Upset plot representing number of significant differentially abundant features from each of the pairwise comparisons with mock treatment after treatment with *A. alternata,* chitin and *A. solani* at 3 and 24 hours post inoculation and the numbers of overlapping features between each of them (intersection size, y-axis) for positive ionization. (B) differentially abundant features from each of the pairwise comparisons with mock treatment after treatment with *A. alternata*, chitin and *A. solani* at 3 and 24 hours post inoculation and the numbers of overlapping features between each of them (intersection size, y-axis) for negative ionization. (C) Venn diagram showing the shared features between *A. alternata* and chitin treatments at 3 and 24 hours post inoculation for positive and negative ionization.

### Candidate metabolites associated with chitin and A. alternata resistance in S. lycopersicum

To look for specific candidate metabolites that can be pattern-triggered and related to *A. alternata* resistance we filtered the compounds shared between chitin and *A. alternata* treatments at 3 and 24 hpi for positive (34 features) and negative (34 features) ionization (Fig. 4C). A total of 68 candidate features are shared between both treatments and detectable at both times. In most cases, the compounds significantly accumulated at 3 and 24 hpi after treatment with both chitin and *A. alternata* and less abundance was observed in mock and *A. solani* samples. After filtering the features detected in both ionization modes, a total of 65 features were considered as candidate features associated with resistance in *S. lycopersicum* (Sup. Table 1). Most of the candidate features show higher abundance after *A. alternata* and chitin treatments at 3 and 24hpi compared with the lower abundance in *A. solani* and Mock treatments (Fig 5). Of the remaining features, 8 are annotated based on MS1 and MS/MS. These compounds included primary metabolites: amino acids and derivatives, nucleotides and derivatives, sulfur-containing nucleosides, and secondary metabolites: alkaloids, and aromatic acids (Table 1). Some of these compounds are mentioned to be involved in defence in plants. While L-threonine itself is not a direct defence compound, the enzyme threonine deaminase (TD) converts threonine to α-ketobutyrate and ammonia as the committed step in isoleucine (Ile) biosynthesis and contributes to JA responses by producing the Ile needed to make the bioactivate JA-Ile conjugate, important for defence against necrotrophs (Gonzales-Vigil et al., 2011; Yeo et al., 2023). 1-Aminocyclopropanecarboxylic acid (ACC) acts as the direct precursor of the plant hormone ethylene, which regulates plant growth and biotic and abiotic stress responses (Van de Poel & Van Der Straeten, 2014; Zaynab et al., 2018). Trigonelline has been shown to accumulate in non-leguminous plants in response to stress, suggesting that it may play a role in stress adaptation (Tyihák et al., 1988). Additionally, trigonelline has been shown to have antioxidant properties, which may help protect plants from oxidative stress caused by biotic and abiotic stressors. Thus, we find several defence associated compounds that are pattern-triggered and can be associated with *A. alternata* resistance, but not *A. solani* susceptibility.

**Figure 5.**
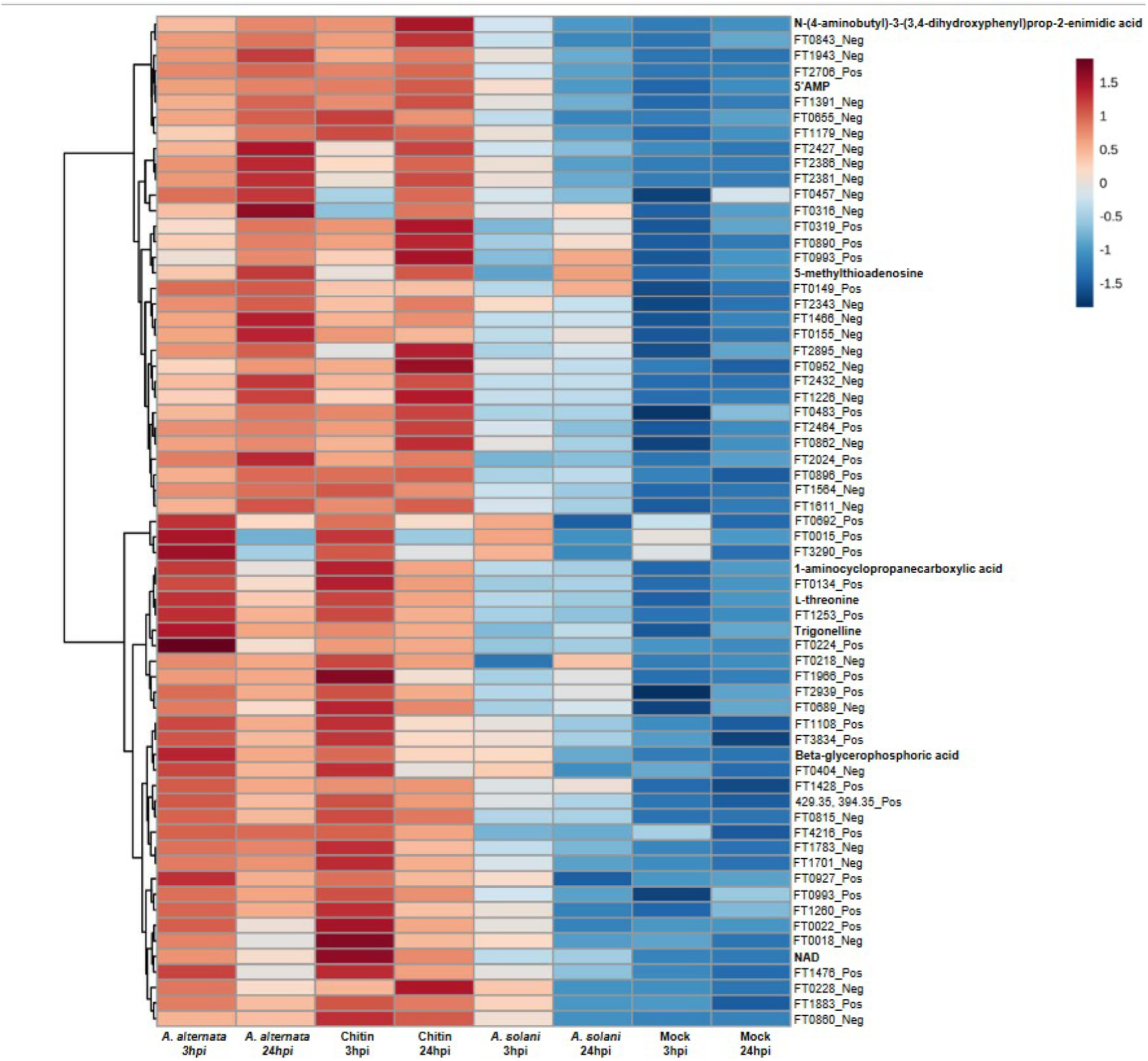
**Heatmap of the total (65) features of interest (positive “_Pos” and negative “_Neg” ionization)** that overlap between A. alternata and chitin treatments at 3 and 24 hours post inoculation.

**Table 1.**
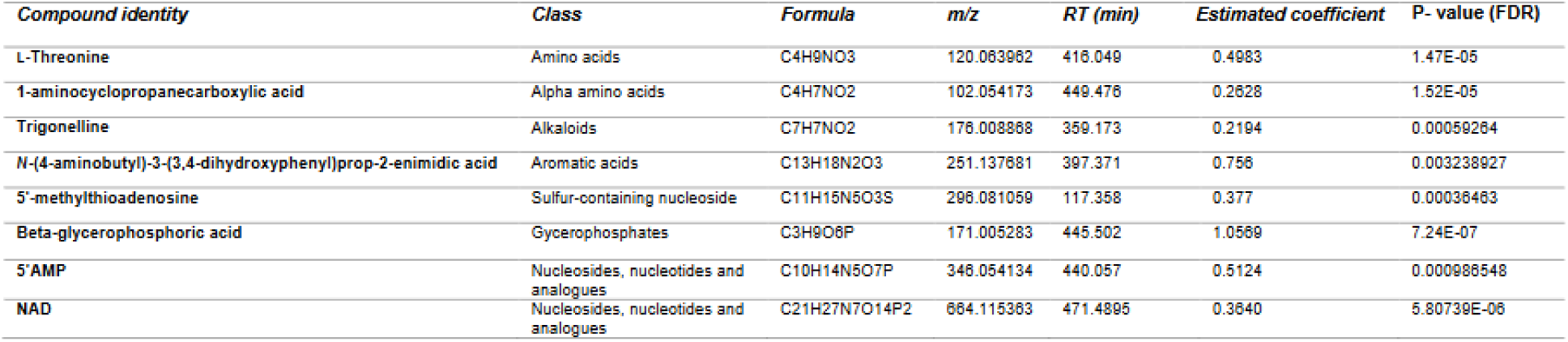
List of candidate metabolites associated to immune response after treatment with *A. alternata* and *A. solani*

### Physiological concentrations of trigonelline show antifungal activity

Because of the assumed high relevance and commercial availability of trigonelline, it was selected for a bioassay to validate its anti-fungal activity effect at physiological concentrations on *A. solani* and *A. alternara* isolates. We observed a significant reduction in the growth of the *A. solani* and *A. alternata* isolates after supplementing SNA growth medium with trigonelline (Fig 6A). After 14 days, the fungal plate growth diameter for both isolates showed clear concentration-dependent reduction. For both isolates, trigonelline concentrations corresponding to the level found in tomato leaves (328 ppm) significantly inhibited fungal growth. Nicotinic acid, a precursor for biosynthesis of trigonelline was also tested and showed marked lower efficacy (Fig 6B). While fungal growth was completely inhibited for concentrations above the physiological levels on the tomato leaves for trigonelline, higher concentrations of nicotinic acid still allowed for some fungal growth. The spores’ germination success was also tested in SNA medium supplemented with trigonelline (Fig 6C). An inhibition of the germination was observed in both fungal cultures at 3.28 mg/mL. Surprisingly, we already observed an inhibition of *A. solani* spores at 0.328 mg/mL, suggesting that trigonnelline is indeed a potent anti-fungal compound.

**Figure 6.**
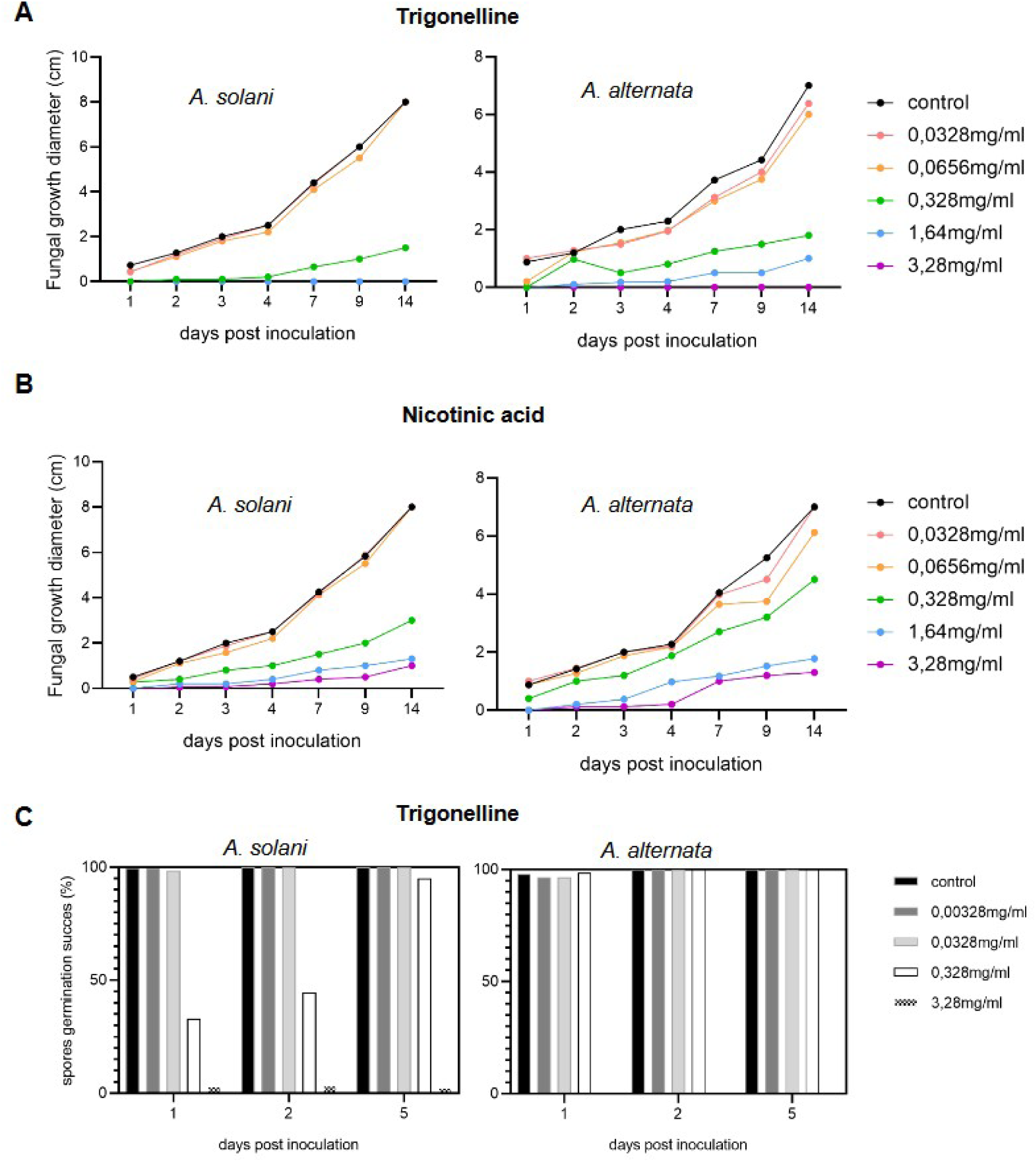
Antifungal activity of trigonelline on Alternaria strains. **(A)** *A. solani* and *A. alternata* growth in cm (y-axis) measured over 14 days after inoculation (x axis) on agar plates using different concentrations of trigonelline. **(B)** *A. solani* and *A. alternata* growth in cm (y-axis) measured over 14 days after inoculation (x axis) on agar plates using different concentrations of nicotinic acid. **(C)** Spore germination success of *A. solani* and *A. alternata* after 1, 2 and 5 days of inoculation using different concentrations of trigonelline in agar plates.

## Discussion

*A. alternata* and *A. solani* are two closely related species of plant pathogens that can cause early blight. While both pathogens can potentially infect tomatoes and potatoes, we identified an *A. alternata* isolate (CS046) that was not able to infect Heinz1706 tomato plants, whereas to *A. solani* (1117-1) caused clear infection symptoms. In order to understand the mechanisms underlying this effect, we set out to determine which metabolic host factors can be associated to *A. alternata* resistance. Tomato plants have developed elaborate defence mechanisms against plant pathogens to survive. These mechanisms include both constitutive and induced defences. Constitutive defences are always present and include physical barriers and potential toxins such as glycoalkaloids (Nakayasu et al., 2021; Zhao et al., 2022). Induced defences are activated in response to pathogen attack and include the production of pathogenesis-related (PR) proteins, phytohormones, and secondary metabolites such as flavonoids and terpenoids (Treutter, 2006; Wink, 2008).

One of the well-known glycoalkaloids produced by tomato plants as defence mechanisms against pathogens is *α*-tomatine (Pegg & Woodward, 1986). This compound is toxic to many pathogens, including fungi, bacteria, viruses, and predatory insects (Duffey & Stout, 1996; Nakayasu et al., 2021). The toxicity mechanism of *α*-tomatine occurs by disrupting cellular membranes by binding to its sterols, leading to membrane rupture and the leakage of cellular contents (Bailly, 2021). Pathogens have evolved mechanisms to detoxify *α*-tomatine, which allows them to infect tomato plants (Oka et al., 2006). We observed that both fungi were able to detoxify *α*-tomatine in tomato broth media, converting this compound into other molecules. Whereas, both isolates were capable of detoxifying *α*-tomatine, it seems that they use different mechanisms for this. It has been described that an active way of dealing with *α*-tomatin is to secrete enzymes that degrade this compound to reduce its toxicity (Osbourn, 1996b; Sandrock & VanEtten, 1998). The degradation process can be categorized into three main actions, based on the hydrolysis products *β*_2_-tomatine, *β*_1_-tomatine and the aglycon tomatidine (You & van Kan, 2021). In our experiment, tomatidine, as well as *β*_1_-tomatine appeared as final breakdown products with higher accumulation in the medium treated with *A. alternata*. Interestingly, (23*R*)-23-acetoxytomatidine accumulated in the medium treated with *A. solani*. (23*R*)-23-acetoxytomatidine was found in the roots of tomato stock and has a fungi-toxic activity, inhibiting the growth of *Fusarium oxysporum* f. sp. *radices-lycopersici* (Nagaoka et al., 1995), yet it appears to not affect *A. solani*. The detected detoxification products of *α*-tomatine explain why both fungi were able to grow in the host broth media and suggest that induced defence responses, rather than phytoanticipins, explain the difference in resistance of the tomato plants against our *A. solani* and *A. alternata isola*tes.

Untargeted metabolomics is a powerful tool that has been used recently for the identification of anti-fungal compounds in plants. For example, one study used untargeted metabolomics to identify steroidal saponins in the foliage and tubers of potato plants as anti-oomycete compounds effective against *Phytophthora infestans* (Baur et al., 2022). Another study used an untargeted metabolite profiling approach on leaf surface of the susceptible cultivated potato *S. tuberosum* and the resistant wild potato species *S. bulbocastanum* to the pathogen *P. infestans*. They found that lysophosphatidylcholine (LPC17:1) was accumulating on the surface of the wild potato, but not on *S. bulbocastanum* and *in vitro* assays revealed an antifungal activity against *P. infestans* (Gorzolka et al., 2021). Secondary metabolites associated with recentance were also focus of another recent study in *S. commersonii;* the differences between accessions that are resistant against *A. solani* and those that are susceptible, can be explained by differences in glycosyltransferase genes that are involved in the production of tetraose steroidal glycoalkaloids that are toxic to *A. solani* (Wolters et al., 2023). In this study, we investigate the potential SMs that may play a role in the resistance of cultivated tomato plants against *A. alternata*. We found a clear reprogramming of metabolomic profiles of tomato leaves after pathogen and chitin treatment. Differential compounds included primary metabolites such as amino acids, lipids and carbohydrates and SMs such as glucosinolates, phenylpropanoids, and organic acids. Although differences existed between metabolomics profiles of *A. alternata* and chitin, many commonly up-regulated metabolites were detected. Small signaling molecules like 1-aminocyclopropane-1-carboxylic acid (ACC) accumulated in samples after the treatments mentioned above. ACC is a precursor of ethylene (ET) which is a plant hormone involved in various physiological processes, including plant defence against necrotrophic pathogens (Zaynab et al., 2018). Enhanced ET production is an early, active response of plants to the perception of pathogen attack and is associated with the induction of disease resistance in plants (Van Loon et al., 2006). ACC has also been proposed to regulate plant development and growth independent of ET, and it can be easily transported over short and long distances, providing the plant with an elaborate system to control local and remote ethylene responses (Van de Poel & Van Der Straeten, 2014). Seeing the involvement of ACC, we suppose that ET signaling plays an important role in resistance against *A. alternata* and that *A. solani* might suppress the ET signaling for a successful infection.

We also observed the accumulation of trigonelline after the treatment with *A. alternata* and chitin. This alkaloid is associated with a number of processes occurring in plants, such as cell cycle regulation, plant growth, and defence (Minorsky, 2002). Trigonelline, which was extracted for the first time from fenugreek seeds, has a well-described biosynthetic pathway and is relatively well-studied for pharmacological activities (Mohamadi et al., 2018). However, there is a limited number of studies that specifically investigate the role of trigonelline in plant defence (Ashihara, 2006; De-la-Cruz Chacón et al., 2013; Sabino et al., 2019).

We validated the role of trigonelline in two bioassays. We show that the addition of this compound to SNA medium resulted in an inhibition of fungal growth of isolates from two *Alternaria* spp. We also tested nicotinic acid, a precursor of trigonelline (Ashihara, 2006), which did not show a complete inhibition of the fungal growth at similar concentrations, thus corroborating the mycotoxic activity of trigonelline. We used concentrations calculated from leaf extract concentrations but also speculate that at the fungal interface, defending cells might accumulate even higher levels of trigonelline.

Our data also suggests that *A. solani* suppresses PTI for a successful infection in tomato plants. The samples with this treatment showed the least amount of compounds accumulated after 24 hpi, and many candidate defence-response compounds, shared between *A. alternata* and chitin treatments, did not significantly accumulate in tomato leaves treated with *A. solani.* To date, the number of reported functional effectors in *A. solani* has been limited and most of them seem to be associated with induction of cell-death (Wang et al., 2022). Besides peptide effectors, *Alternaria* species can synthesize phytotoxic metabolites, such as AAL toxins and other host-specific or host-nonspecific toxins. Such toxins serve chemical inhibition of defence compounds produced by the host or the induction of cell death (Wang et al., 2023). There is limited information on the specific role of toxins of *A. solani* in pathogen infection. However, a recent study showed the effect of alternaric acid, a toxin of *A. solani*, on the hypersensitive response of potato to *P. infestans* (Wang et al., 2023). Besides deeper investigations in other host defence mechanisms, follow-up studies on the identification of effectors or possible other small molecules that are responsible for PTI suppression by *A. solani* during infection of cultivated tomato plants could be of interest for further understanding of this host-pathogen interactions.

Overall, our findings show the presence of pattern-triggered chemical defence barriers against *A. alternata*, which likely contribute to the resistance of cultivated tomato plants. They also illustrate how untargeted metabolomics can help in the elucidation of new antifungal compounds that can may be of interest for future crop protection strategies.

## Acknowledgments

We acknowledge Nicole Bellé (Chair of Phytopathology, TUM) for providing *A. solani* (1117-1) isolate. This work was supported by the German Research Foundation (SFB924, projects: B8, B12 and B13).

---

**Supplementary Figure 1.**
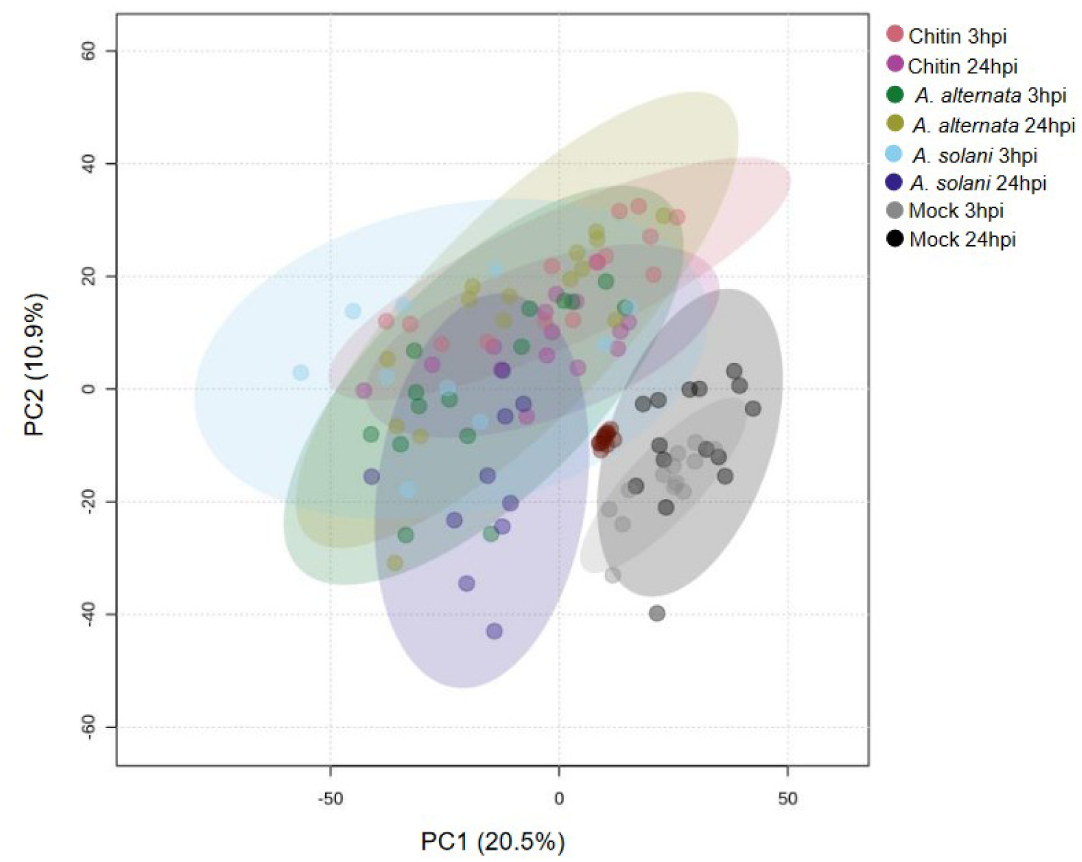
PCA representation of the overview of the metabolomic profiles from 3 different treatments (*A. alternata*, chitin and *A. solani*) compared with mock samples (treated with water) after 3 and 24 hours post inoculation. (*n*=15). Confidence regions of 95% marked in circles for each group. Axes represent the first (x axis) and second (y axis) principal component, the percentage of variance explained is shown on each axis.

**Supplementary Figure 2.**
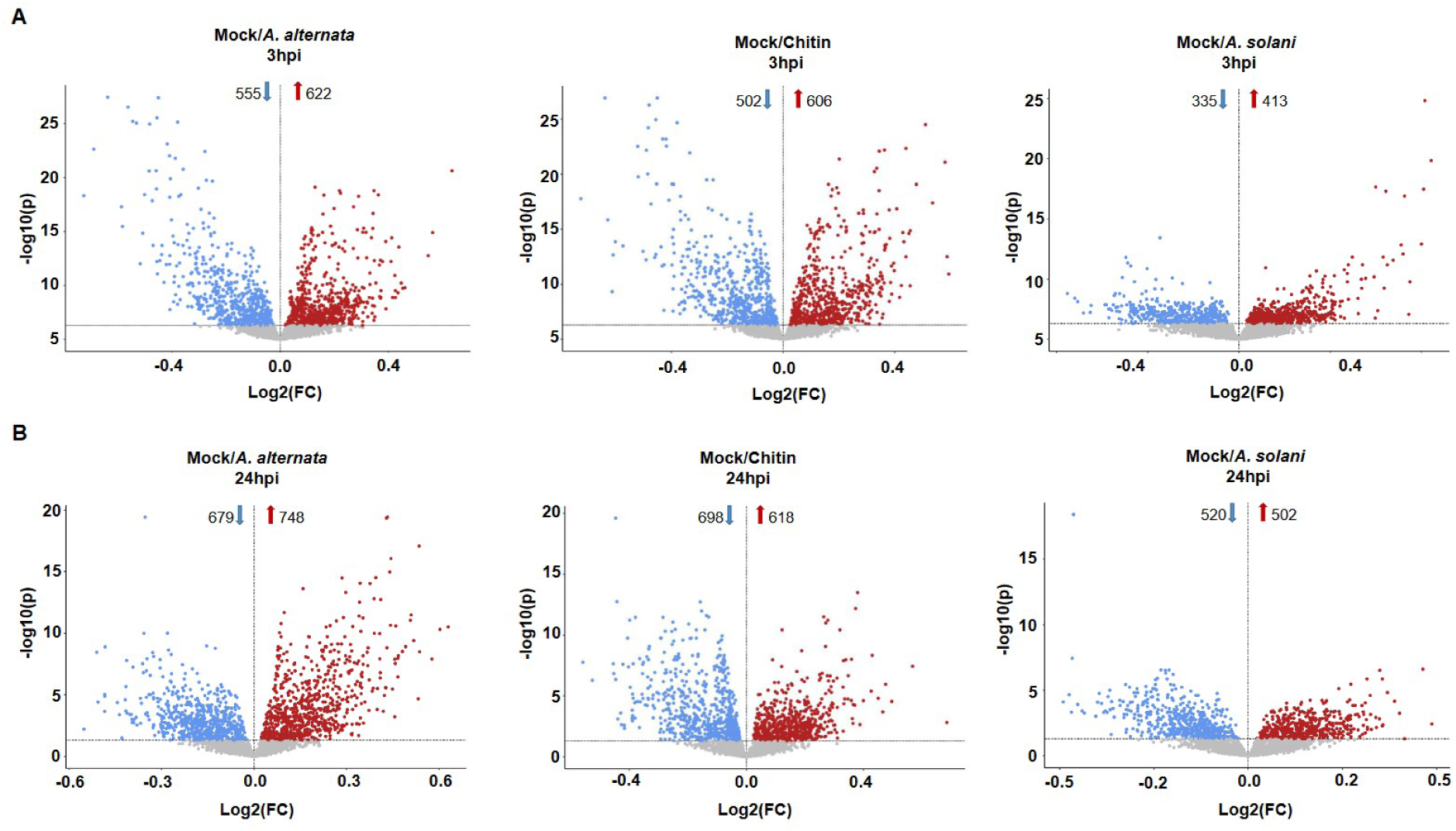
(**A)** Volcano plots showing the overview of the significant features for *A. alternata,* chitin *and A, solani* vs. mock treatments after 3 hours of inoculation. **(B)** Volcano plots showing the overview of the significant features for *A. alternata,* chitin *and A, solani* vs. mock treatments after 24 hours of inoculation. Red and blue color represent the increased and decreased accumulation of metabolites, respectively. X-axis showing Log2(FC) y-axis showing – log10 (p) (*P*-value threshold < 0.05 with FDR and FC >1).

**Supplementary Figure 3.**
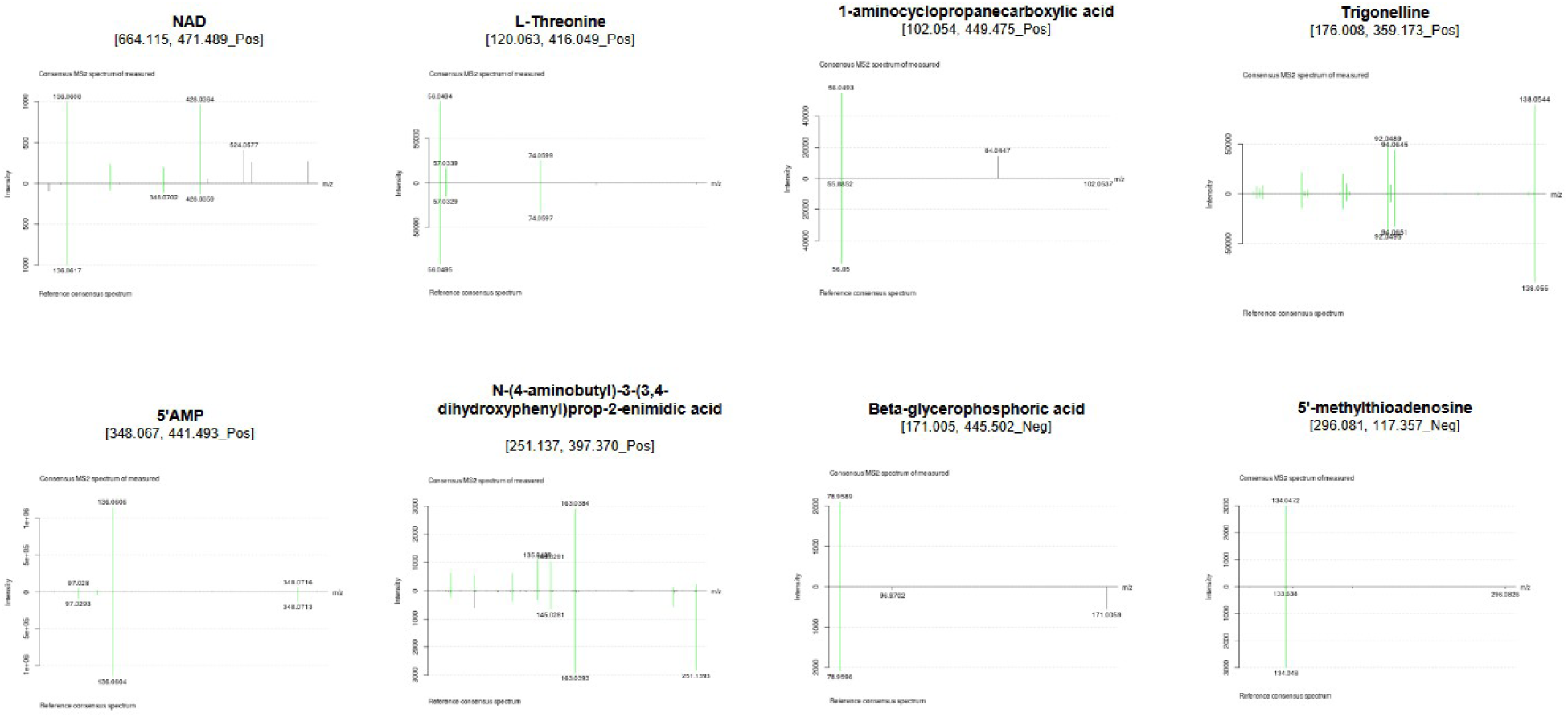
MS2 annotation spectra of significantly abundant metabolic features after treatment with *A. alternata* and chitin at 3 and 24hpi

## Notes

### Competing Interest Statement

The authors have declared no competing interest.

## References

Adhikari, P., Oh, Y., & Panthee, D. R. (2017). Current Status of Early Blight Resistance in Tomato: An Update. International Journal of Molecular Sciences, 18(10), Article 10. https://doi.org/10.3390/ijms18102019

Ahmad, F., Raziq, F., Ullah, N., Khan, H., & Din, N. (2017). In vitro and in vivo bio-assay of phytobiocidal effect of plant extracts on Alternaria solani causing agent of early blight disease in tomato. Archives of Phytopathology and Plant Protection, 50(11–12), 568–583. https://doi.org/10.1080/03235408.2017.1352247

Akhtar, K. P., Ullah, N., Saleem, M. Y., Iqbal, Q., Asghar, M., & Khan, A. R. (2019). Evaluation of tomato genotypes for early blight disease resistance caused by Alternaria solani in Pakistan. Journal of Plant Pathology, 101(4), 1159–1170.

Allwood, J. W., Williams, A., Uthe, H., van Dam, N. M., Mur, L. A. J., Grant, M. R., & Pétriacq, P. (2021). Unravelling Plant Responses to Stress—The Importance of Targeted and Untargeted Metabolomics. Metabolites, 11(8), 558. https://doi.org/10.3390/metabo11080558

Ashihara, H. (2006). Metabolism of alkaloids in coffee plants. Brazilian Journal of Plant Physiology, 18, 1–8. https://doi.org/10.1590/S1677-04202006000100001

Bailly, C. (2021). The steroidal alkaloids α-tomatine and tomatidine: Panorama of their mode of action and pharmacological properties. Steroids, 176, 108933. https://doi.org/10.1016/j.steroids.2021.108933

Baur, S., Bellé, N., Frank, O., Wurzer, S., Pieczonka, S. A., Fromme, T., Stam, R., Hausladen, H., Hofmann, T., Hückelhoven, R., & Dawid, C. (2022). Steroidal Saponins─New Sources to Develop Potato (Solanum tuberosum L.) Genotypes Resistant against Certain Phytophthora infestans Strains. Journal of Agricultural and Food Chemistry, 70(24), 7447– 7459. https://doi.org/10.1021/acs.jafc.2c02575

Benton, H. P., Want, E. J., & Ebbels, T. M. D. (2010). Correction of mass calibration gaps in liquid chromatography–mass spectrometry metabolomics data. Bioinformatics, 26(19), 2488– 2489. https://doi.org/10.1093/bioinformatics/btq441

Bozbuga, R., Ates, S. Y., Guler, P. G., Yildiz, H. N., Kara, P. A., Arpaci, B. B., Imren, M., Bozbuga, R., Ates, S. Y., Guler, P. G., Yildiz, H. N., Kara, P. A., Arpaci, B. B., & Imren, M. (2022). Host-Pathogen and Pest Interactions: Virus, Nematode, Viroid, Bacteria, and Pests in Tomato Cultivation. In Tomato—From Cultivation to Processing Technology. IntechOpen. https://doi.org/10.5772/intechopen.106064

Chaerani, R., Groenwold, R., Stam, P., & Voorrips, R. E. (2007). Assessment of early blight (Alternaria solani) resistance in tomato using a droplet inoculation method. Journal of General Plant Pathology, 73(2), 96–103. https://doi.org/10.1007/s10327-006-0337-1

Chaerani, R., & Voorrips, R. E. (2006). Tomato early blight (Alternaria solani): The pathogen, genetics, and breeding for resistance. Journal of General Plant Pathology, 72(6), 335–347. https://doi.org/10.1007/s10327-006-0299-3

De-la-Cruz Chacón, I., Riley-Saldaña, C. A., & González-Esquinca, A. R. (2013). Secondary metabolites during early development in plants. Phytochemistry Reviews, 12(1), 47–64. https://doi.org/10.1007/s11101-012-9250-8

Duffey, S. S., & Stout, M. J. (1996). Antinutritive and toxic components of plant defense against insects. Archives of Insect Biochemistry and Physiology, 32(1), 3–37. https://doi.org/10.1002/(SICI)1520-6327(1996)32:1<3::AID-ARCH2>3.0.CO;2-1

Dührkop, K., Fleischauer, M., Ludwig, M., Aksenov, A. A., Melnik, A. V., Meusel, M., Dorrestein, P. C., Rousu, J., & Böcker, S. (2019). SIRIUS 4: A rapid tool for turning tandem mass spectra into metabolite structure information. Nature Methods, 16(4), Article 4. https://doi.org/10.1038/s41592-019-0344-8

FAOSTAT. Statistical Database. (2020). Food and Agriculture Organization of the United Nations, Rome. [Accessed:12/05/2023, 17:52:12]

Glas, J., Schimmel, B., Alba, J., Escobar-Bravo, R., Schuurink, R., & Kant, M. (2012). Plant Glandular Trichomes as Targets for Breeding or Engineering of Resistance to Herbivores. International Journal of Molecular Sciences, 13(12), 17077–17103. https://doi.org/10.3390/ijms131217077

Gonzales-Vigil, E., Bianchetti, C. M., Phillips, G. N., & Howe, G. A. (2011). Adaptive evolution of threonine deaminase in plant defense against insect herbivores. Proceedings of the National Academy of Sciences, 108(14), 5897–5902. https://doi.org/10.1073/pnas.1016157108

Gorzolka, K., Perino, E. H. B., Lederer, S., Smolka, U., & Rosahl, S. (2021). Lysophosphatidylcholine 17:1 from the Leaf Surface of the Wild Potato Species Solanum bulbocastanum Inhibits Phytophthora infestans. Journal of Agricultural and Food Chemistry, 69(20), 5607–5617. https://doi.org/10.1021/acs.jafc.0c07199

Landeo Villanueva, S., Malvestiti, M. C., van Ieperen, W., Joosten, M. H. A. J., & van Kan, J. A. L. (2021). Red light imaging for programmed cell death visualization and quantification in plant–pathogen interactions. Molecular Plant Pathology, 22(3), 361–372. https://doi.org/10.1111/mpp.13027

Lê Cao, K.-A., Boitard, S., & Besse, P. (2011). Sparse PLS discriminant analysis: Biologically relevant feature selection and graphical displays for multiclass problems. BMC Bioinformatics, 12(1), 253. https://doi.org/10.1186/1471-2105-12-253

Minorsky, P. V. (2002). The Hot and the Classic. Plant Physiology, 128(1), 7–8. https://doi.org/10.1104/pp.900014

Moghaddam, G., Nasr Esfahani, M., Rezayatmand, Z., & Khozaei, M. (2022). Genomic markers analysis associated with resistance to Alternaria alternata (fr.) keissler-tomato pathotype, Solanum lycopersicum L. Breeding Science, 72. https://doi.org/10.1270/jsbbs.22003

Mohamadi, N., Sharififar, F., Pournamdari, M., & Ansari, M. (2018). A Review on Biosynthesis, Analytical Techniques, and Pharmacological Activities of Trigonelline as a Plant Alkaloid. Journal of Dietary Supplements, 15(2), 207–222. https://doi.org/10.1080/19390211.2017.1329244

Muñoz-Hoyos, L., & Stam, R. (2023). Metabolomics in plant pathogen-defence: From single molecules to large scale analysis. Phytopathology. https://doi.org/10.1094/PHYTO-11-22-0415-FI

Nagaoka, T., Ohra, J., Yoshihara, T., & Sakamura, S. (1995). Fungitoxic Compounds from the Roots of Tomato Stock. Japanese Journal of Phytopathology, 61(2), 103–108. https://doi.org/10.3186/jjphytopath.61.103

Nakayasu, M., Ohno, K., Takamatsu, K., Aoki, Y., Yamazaki, S., Takase, H., Shoji, T., Yazaki, K., & Sugiyama, A. (2021). Tomato roots secrete tomatine to modulate the bacterial assemblage of the rhizosphere. Plant Physiology, 186(1), 270–284. https://doi.org/10.1093/plphys/kiab069

Oka, K., Okubo, A., Kodama, M., & Otani, H. (2006). Detoxification of α-tomatine by tomato pathogens Alternaria alternata tomato pathotype and Corynespora cassiicola and its role in infection. Journal of General Plant Pathology, 72(3), 152–158. https://doi.org/10.1007/s10327-005-0262-8

Ökmen, B., Etalo, D. W., Joosten, M. H. A. J., Bouwmeester, H. J., de Vos, R. C. H., Collemare, J., & de Wit, P. J. G. M. (2013). Detoxification of α-tomatine by Cladosporium fulvum is required for full virulence on tomato. New Phytologist, 198(4), 1203–1214. https://doi.org/10.1111/nph.12208

Osbourn, A. (1996a). Preformed Antimicrobial Compounds and Plant Defense against Fungal Attack. The Plant Cell, 8(10), 1821–1831.

Osbourn, A. (1996b). Saponins and plant defence—A soap story. Trends in Plant Science, 1(1), 4–9. https://doi.org/10.1016/S1360-1385(96)80016-1

Özer Çalış. (2011). Genetic analysis of resistance to early blight disease in tomato. AFRICAN JOURNAL OF BIOTECHNOLOGY, 10(79). https://doi.org/10.5897/AJB11.1872

Pandey, D., Rajendran, S. R. C. K., Gaur, M., Sajeesh, P. K., & Kumar, A. (2016). Plant Defense Signaling and Responses Against Necrotrophic Fungal Pathogens. Journal of Plant Growth Regulation, 35(4), 1159–1174. https://doi.org/10.1007/s00344-016-9600-7

Pegg, G. F., & Woodward, S. (1986). Synthesis and metabolism of α-tomatine in tomato isolines in relation to resistance to Verticillium albo-atrum. Physiological and Molecular Plant Pathology, 28(2), 187–201. https://doi.org/10.1016/S0048-4059(86)80063-7

Quidde, T., Osbourn, A. E., & Tudzynski, P. (1998). Detoxification of α-tomatine byBotrytis cinerea. Physiological and Molecular Plant Pathology, 52(3), 151–165. https://doi.org/10.1006/pmpp.1998.0142

Ray, S., Mondal, S., Chowdhury, S., & Kundu, S. (2015). Differential responses of resistant and susceptible tomato varieties to inoculation with Alternaria solani. Physiological and Molecular Plant Pathology, 90, 78–88. https://doi.org/10.1016/j.pmpp.2015.04.002

Roddick, J. G. (1974). The steroidal glycoalkaloid α-tomatine. Phytochemistry, 13(1), 9–25. https://doi.org/10.1016/S0031-9422(00)91261-5

Roldán-Arjona, T., Pérez-Espinosa, A., & Ruiz-Rubio, M. (1999). Tomatinase from Fusarium oxysporum f. Sp. Lycopersici Defines a New Class of Saponinases. Molecular Plant-Microbe Interactions®, 12(10), 852–861. https://doi.org/10.1094/MPMI.1999.12.10.852

Sabino, A. R., Tavares, S. S., Riffel, A., Li, J. V., Oliveira, D. J. A., Feres, C. I. M. A., Henrique, L., Oliveira, J. S., Correia, G. D. S., Sabino, A. R., Nascimento, T. G., Hawkes, G., Santana, A. E. G., Holmes, E., & Bento, E. S. (2019). 1H NMR metabolomic approach reveals chlorogenic acid as a response of sugarcane induced by exposure to Diatraea saccharalis. Industrial Crops and Products, 140, 111651. https://doi.org/10.1016/j.indcrop.2019.111651

Sadeghi, B., Mirzaei, S., & Fatehi, F. (2022). The proteomic analysis of the resistance responses in tomato during interaction with Alternaria alternate. Scientia Horticulturae, 304, 111295. https://doi.org/10.1016/j.scienta.2022.111295

Sandrock, R. W., & VanEtten, H. D. (1998). Fungal Sensitivity to and Enzymatic Degradation of the Phytoanticipin α-Tomatine. Phytopathology®, 88(2), 137–143. https://doi.org/10.1094/PHYTO.1998.88.2.137

Schauer, N., Semel, Y., Balbo, I., Steinfath, M., Repsilber, D., Selbig, J., Pleban, T., Zamir, D., & Fernie, A. R. (2008). Mode of Inheritance of Primary Metabolic Traits in Tomato. The Plant Cell, 20(3), 509–523. https://doi.org/10.1105/tpc.107.056523

Schmey, T., Small, C., Hoyoz, L. M., Ali, T., Gamboa, S., Mamami, B., Sepulveda, G. C., Thines, M., & Stam, R. (2022). Small-spored Alternaria spp. (Section Alternaria) are common pathogens on wild tomato species [Preprint]. Microbiology. https://doi.org/10.1101/2022.12.08.519636

Simmons, E. G. (2007). Alternaria: An Identification Manual: Fully Illustrated and with Catalogue Raisonné 1796-2007. CBS Fungal Biodiversity Centre. https://books.google.co.uk/books?id=F\_AzKgAACAAJ

Smith, C. A., Want, E. J., O’Maille, G., Abagyan, R., & Siuzdak, G. (2006). XCMS: Processing Mass Spectrometry Data for Metabolite Profiling Using Nonlinear Peak Alignment, Matching, and Identification. Analytical Chemistry, 78(3), 779–787. https://doi.org/10.1021/ac051437y

Taiz, L., & Zeiger, E. (1991). Plant Physiology. Benjamin/Cummings Publishing Company. https://books.google.co.uk/books?id=KeIUAQAAIAAJ

Tautenhahn, R., Böttcher, C., & Neumann, S. (2008). Highly sensitive feature detection for high resolution LC/MS. BMC Bioinformatics, 9(1), 504. https://doi.org/10.1186/1471-2105-9-504

Treutter, D. (2006). Significance of flavonoids in plant resistance: A review. Environmental Chemistry Letters, 4(3), 147–157. https://doi.org/10.1007/s10311-006-0068-8

Tyihák, E., Sarhan, A. R. T., Cong, N. T., Barna, B., & Király, Z. (1988). The level of trigonelline and other quaternary ammonium compounds in tomato leaves in ratio to the changing nitrogen supply. Plant and Soil, 109(2), 285–287. https://doi.org/10.1007/BF02202097

Van de Poel, B., & Van Der Straeten, D. (2014). 1-aminocyclopropane-1-carboxylic acid (ACC) in plants: More than just the precursor of ethylene! Frontiers in Plant Science, 5. https://www.frontiersin.org/articles/10.3389/fpls.2014.00640

Van Loon, L. C., Geraats, B. P. J., & Linthorst, H. J. M. (2006). Ethylene as a modulator of disease resistance in plants. Trends in Plant Science, 11(4), 184–191. https://doi.org/10.1016/j.tplants.2006.02.005

Wang, C., Wang, J., Zhang, D., Cheng, J., Zhu, J., & Yang, Z. (2023). Identification and functional analysis of protein secreted by Alternaria solani. PLOS ONE, 18(3), e0281530. https://doi.org/10.1371/journal.pone.0281530

Wang, C., Zhang, D., Cheng, J., Zhao, D., Pan, Y., Li, Q., Zhu, J., Yang, Z., & Wang, J. (2022). Identification of effector CEP112 that promotes the infection of necrotrophic Alternaria solani. BMC Plant Biology, 22(1), 466. https://doi.org/10.1186/s12870-022-03845-w

Wink, M. (2008). Plant Secondary Metabolism: Diversity, Function and its Evolution. Natural Product Communications, 3(8), 1934578X0800300. https://doi.org/10.1177/1934578X0800300801

Wishart, D. S., Tzur, D., Knox, C., Eisner, R., Guo, A. C., Young, N., Cheng, D., Jewell, K., Arndt, D., Sawhney, S., Fung, C., Nikolai, L., Lewis, M., Coutouly, M.-A., Forsythe, I., Tang, P., Shrivastava, S., Jeroncic, K., Stothard, P.,… Querengesser, L. (2007). HMDB: The Human Metabolome Database. Nucleic Acids Research, 35(Database issue), D521–526. https://doi.org/10.1093/nar/gkl923

Wolters, Doret, W., M, T. Y., Shimlal, A., P, K. L., Miriam, S., Lotte, C., F, V. R. G., & A, V. V. G. A. (2023). Tetraose steroidal glycoalkaloids from potato can provide complete protection against fungi and insects. ELife, 12. https://doi.org/10.7554/eLife.87135

Xia, J., Psychogios, N., Young, N., & Wishart, D. S. (2009). MetaboAnalyst: A web server for metabolomic data analysis and interpretation. Nucleic Acids Research, 37(suppl_2), W652–W660. https://doi.org/10.1093/nar/gkp356

Yeo, I.-C., de Azevedo Manhaes, A. M. E., Liu, J., Avila, J., He, P., & Devarenne, T. P. (2023). An unexpected role for tomato threonine deaminase 2 in host defense against bacterial infection. Plant Physiology, 192(1), 527–545. https://doi.org/10.1093/plphys/kiac584

You, Y., & van Kan, J. A. L. (2021). Bitter and sweet make tomato hard to (b)eat. New Phytologist, 230(1), 90–100. https://doi.org/10.1111/nph.17104

Zaynab, M., Fatima, M., Abbas, S., Sharif, Y., Umair, M., Zafar, M. H., & Bahadar, K. (2018). Role of secondary metabolites in plant defense against pathogens. Microbial Pathogenesis, 124, 198–202. https://doi.org/10.1016/j.micpath.2018.08.034

Zhao, T., Pei, T., Jiang, J., Yang, H., Zhang, H., Li, J., & Xu, X. (2022). Understanding the mechanisms of resistance to tomato leaf mold: A review. Horticultural Plant Journal, 8(6), 667–675. https://doi.org/10.1016/j.hpj.2022.04.008

